# Beta Diversity Meta-Analysis Shows Transformations Have Broadly Similar Performance in Machine Learning Applications Regardless of Compositional or Phylogenetic Awareness

**DOI:** 10.64898/2026.01.20.699043

**Authors:** Daisy Fry Brumit, Alicia A. Sorgen, Anthony Fodor

## Abstract

**Background:** Beta diversity quantifies pairwise differences between two or more communities through matrix transformations, which are either naïve to phylogeny or phylogenetically aware. Methods have recently been introduced that also consider compositionality and sparsity and that display an increased magnitude of pseudo-F scores as produced by PERMANOVA to measure effect size. In this study, we ask how transformations that consider phylogeny, sparsity, and compositionality compare to older, simpler methods across five publicly available datasets.

**Results:** Application of random forest methods to 107 features across 5 datasets did not yield a consistent increase in classification performance between different beta diversity methods. Limiting datasets to just three eigenvalue decomposition (EVD) axes leads to a small but reliably detectable decrease in performance compared to giving random forest models access to log-normalized or even un-normalized raw count tables. Increasing the number of included EVD axes in classification improves performance across all available models up to ∼10-20 axes. We observed larger variation in PERMANOVA pseudo-F scores for some features associated with phylogenetically and compositionally aware beta diversity algorithms across multiple datasets, but did not find that these improved scores yielded consistently increased resolution or accuracy for machine learning methods.

**Conclusions:** While EVD remains an essential technique for dimension reduction, retaining higher-dimensional structures past 3 EVD axes may improve performance. Elevated but insignificant pseudo-F scores may be explained by the higher variance in pseudo-F scores for phylogenetically or compositionally aware methods compared to simpler methods.This indicates that pseudo-F scores are an unreliable overall metric of algorithm performance. Taken together, our results show that choice of beta diversity metric does not yield a substantial difference in effect size or machine learning performance. We conclude that analysts are free to choose appropriate methods for each dataset balancing simplicity vs. corrections for phylogeny, sparsity and compositionality and that these choices are unlikely to impact the overall power and resolution of biological conclusions from microbial data.

## Background and Introduction

Beta diversity refers to measures of distance between ecological communities and can be quantified a variety of ways through different transformations. The earliest beta diversity measures used in the metagenomics literature were insensitive to phylogeny and compositionality; these included the Jaccard and Bray-Curtis metrics that simply compare communities based on the presence/absence alone and with abundance of individuals, respectively [1,2]. In 2005 and 2007, two forms of the UniFrac algorithm were introduced in a pair of papers that argued incorporating phylogenetic information into beta diversity measures can lead to an enhanced understanding of how microbial communities are structured in relationship to environmental factors [3,4]. Of these two, unweighted UniFrac considers only presence/absence data from communities, while weighted UniFrac incorporates proportions. While popular, the application of UniFrac has not been un-controversial, with some exchanges between the original authors and those who challenge the algorithm’s robustness [5–7].

In addition to phylogeny, researchers are often concerned about the implications of compositionality throughout metagenomic pipelines and analyses, including beta diversity. Compositionality in metagenomics refers to the tendency of metagenomic read counts data to be constrained to sum to an arbitrary value, which violates assumptions present in many canonical statistical analyses. The consequences of this have been the subject of a good deal of attention in the metagenomic literature in ways that are closely tied to concepts of beta diversity [8]. One way of dealing with compositionality is to break count data into proportional space by applying log ratio transformations, such as the centered log ratio (CLR) transformation [9]. Aitchison distance, which was developed in concert with CLR, is one beta diversity metric that is meant to resolve the compositionality issue and is examined in a previous paper [10]. In 2019, Martino et al. proposed a new beta diversity metric called robust principal component analysis (RPCA), which employs CLR in a way that is meant to be more robust to data sparsity.

In 2022, Martino et al. then proposed a new adaptation of this method that also employs the phylogenetic awareness of UniFrac, called phylo-RPCA [11]. This paper examined a cross-sectional study on indoor microbiomes [12], as well as simulated data and a repeated-measures dataset for a complementary model. They applied various forms of UniFrac, Jaccard, Bray-Curts, Aitchison, and their own transformations to the appropriate datasets, output the first three axes of each transformation after eigenvalue decomposition (EVD), and compared the effects of each transformation on resolving phenotype via PERMANOVA and

*k*-nearest neighbors (KNN) classification. These results showed a sizable increase in both pseudo-F values from PERMANOVA and classification performance from KNN, in favor of both phylo-RPCA and its complement, phylo-CTF (compositional tensor factorization). Here, PERMANOVA serves to relate metadata features back to beta diversity transformed tables. While p-values convey significance, pseudo-F values generated by PERMANOVA quantify the ratio between between-group variation and within-group variation when handling multivariate data. Thus, higher pseudo-F values are considered to indicate a better distinction between groups [13].

While impressive, these results were narrowly applied to a small number of datasets using a very specific bioinformatic pipeline. In this paper, we consider whether these previously reported observations can extend to additional datasets and a broader analysis pipeline. Specifically, we used five publicly available 16S rRNA datasets on various topics related to human microbiomes, as seen in a previous meta-analysis and the phylo-RPCA publication [10,11]. While we used PERMANOVA and KNN, we also included a random forest analysis for mapping features back to count data. Random forest is commonly used in metagenomics studies, as it is not dependent on appropriate *k* selection. We also evaluated whether the inclusion of additional dimensions beyond the first three axes or use of all available count data improved classification performance. Our results indicate that these choices involving how to apply beta diversity pipelines showed little overall impact on the performance of our selected ML methods.

## Methods

This meta-analysis aimed to determine whether increasing complexity, phylogenetic awareness, or correction for compositionality and sparsity affect phenotypic resolution in the context of beta diversity models. All scripts associated with this study can be viewed at https://github.com/DaisyBrumit/beta_diversity_testing.

### Datasets

Five publicly available datasets were used in this analysis, whose study information can be found summarized in Table 1. Of these, the Ruiz-Calderon urbanization dataset [12] was used in the original paper by Martino et al. [11]., and the remaining four were used in a previous publication [10]. All five datasets contain 16S rRNA sequences associated with human gut microbiota. In addition to human stool samples, the Jones dataset includes samples taken from mucosal swabs, and the Ruiz-Calderon dataset includes skin microbiome samples.

**Table 1.**
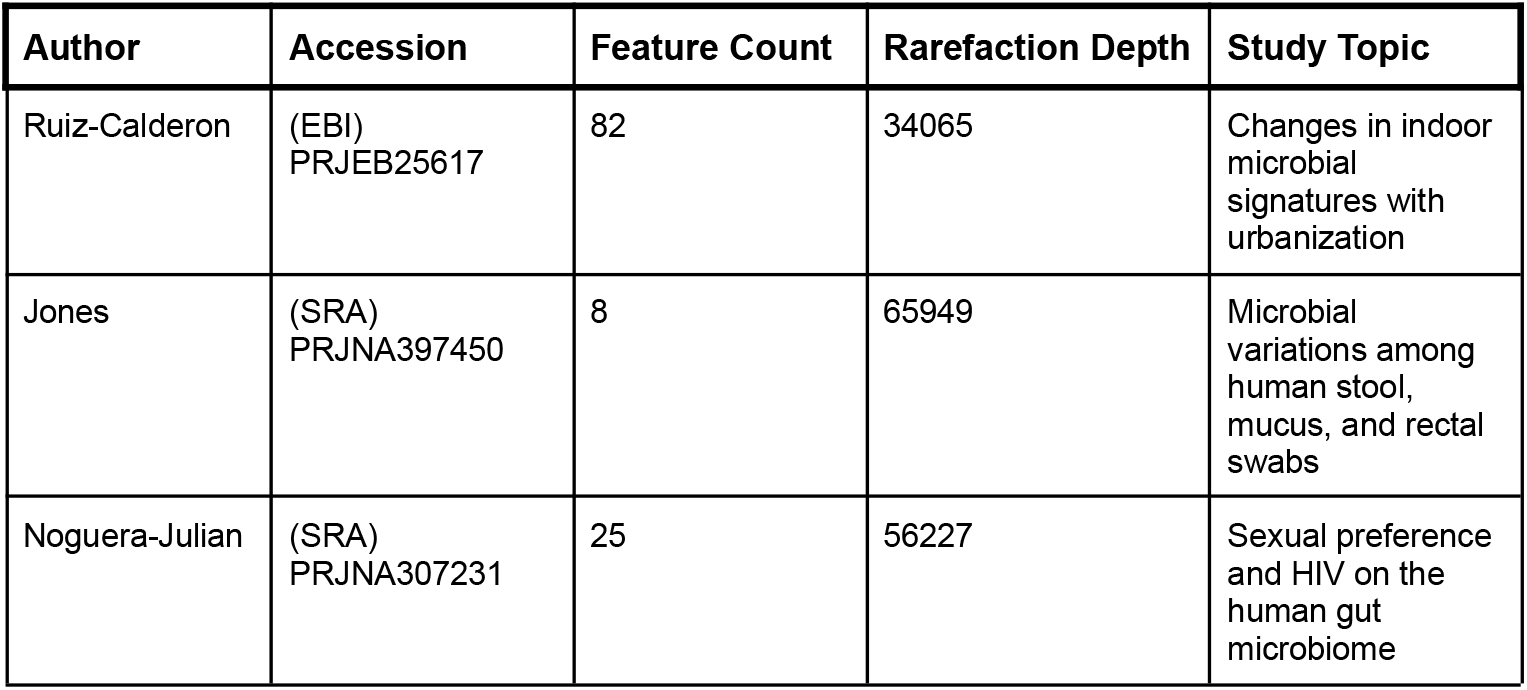
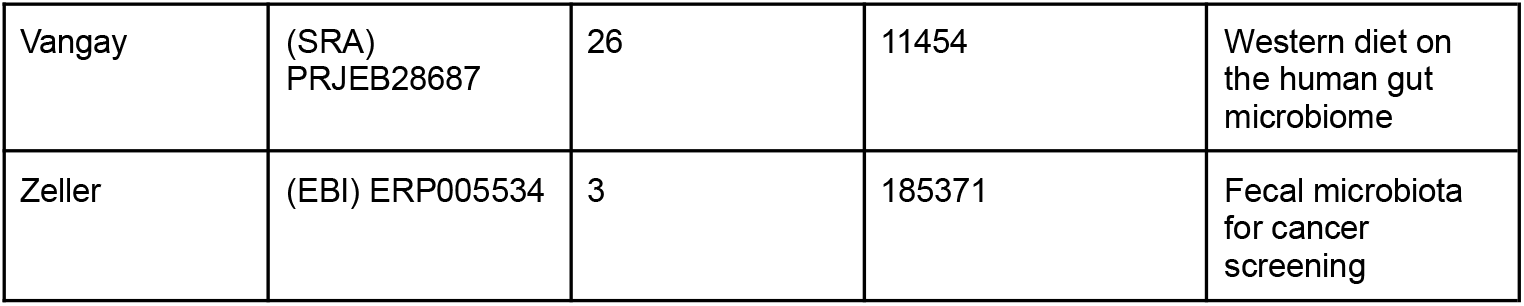
A summary of datasets included for study.

| Author | Accession | Feature Count | Rarefaction Depth | Study Topic |
| --- | --- | --- | --- | --- |
| Ruiz-Calderon | (EBI)<br>PRJEB25617 | 82 | 34065 | Changes in indoor microbial signatures with urbanization |
| Jones | (SRA)<br>PRJNA397450 | 8 | 65949 | Microbial variations among human stool, mucus, and rectal swabs |
| Noguera-Julian | (SRA)<br>PRJNA307231 | 25 | 56227 | Sexual preference and HIV on the human gut microbiome |
| Vangay | (SRA)<br>PRJEB28687 | 26 | 11454 | Western diet on<br>the human gut<br>microbiome |
| Zeller | (EBI) ERP005534 | 3 | 185371 | Fecal microbiota<br>for cancer<br>screening |

### Sequences and Phylogeny

Preprocessing of sequences with DADA2 was conducted previously for four of five studies [10]. To ensure methodological consistency, DADA2 was re-applied to include the Ruiz-Calderon dataset. All taxonomic profiling of 16S rRNA data was conducted with the Greengenes 13-8 reference database [14] for the sake of consistency and compatibility with the Qiime2 framework[15,16]. The reference database was used to create insertion trees in lieu of generating *de novo* trees, and the resulting filtered feature tables were used for downstream analyses.

### Beta Diversity Transformations

After tree building, normalization schemes were applied to the data prior to beta diversity transformation, as detailed in Table 2. For compositionally and phylogenetically naïve transformations (Jaccard, Bray Curtis), the following log normalization, which we call “Lognorm” was applied.

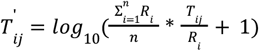

**Table 2.**
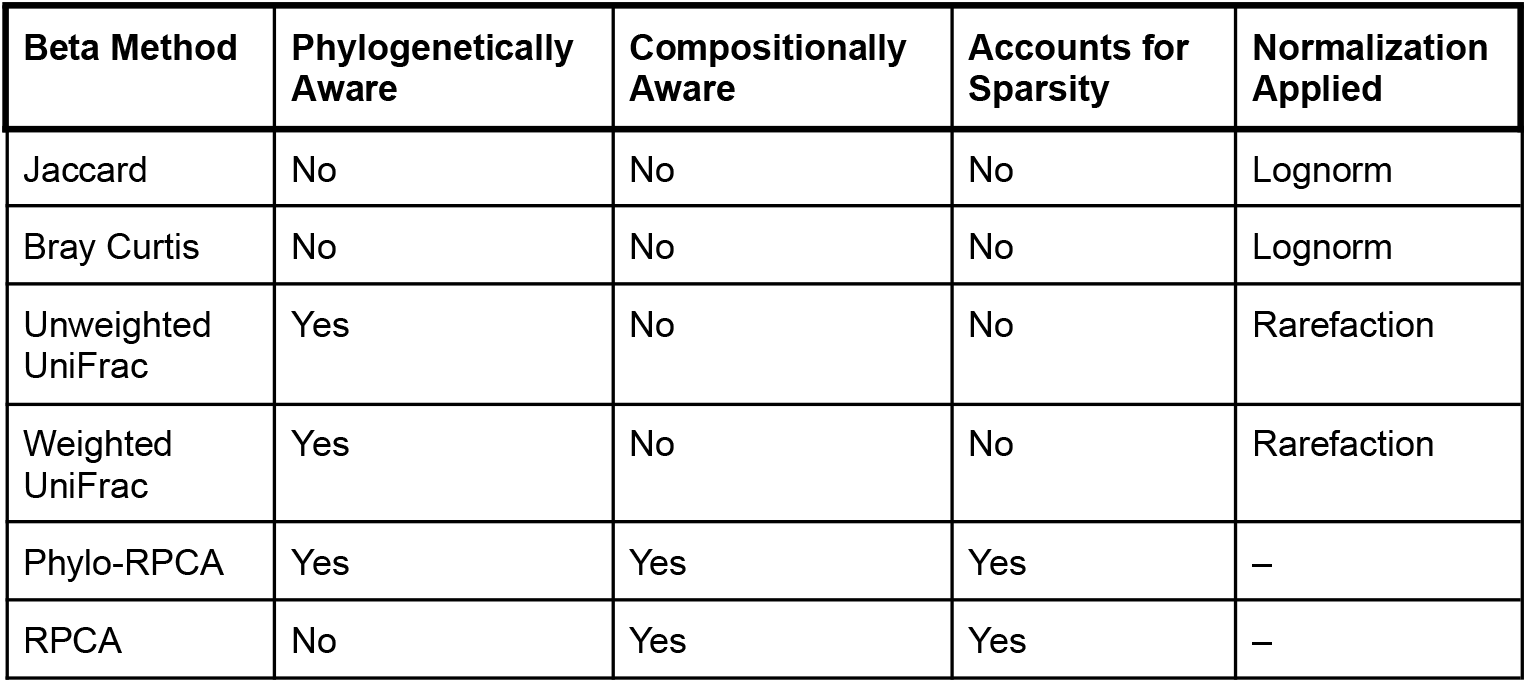
Beta diversity transformations differ across the board in their approach to dealing with statistical assumptions. Normalization schemes will vary based on the beta diversity approach.

| Beta Method | Phylogenetically Aware | Compositionally Aware | Accounts for Sparsity | Normalization Applied |
| --- | --- | --- | --- | --- |
| Jaccard | No | No | No | Lognorm |
| Bray Curtis | No | No | No | Lognorm |
| Unweighted UniFrac | Yes | No | No | Rarefaction |
| Weighted UniFrac | Yes | No | No | Rarefaction |
| Phylo-RPCA | Yes | Yes | Yes | – |
| RPCA | No | Yes | Yes | – |

Where T’_ij_ is the log normalized table, R_i_ is the sum of the i^th^ row, n is the total number of rows, and T_ij_ is a single observation at the i^th^ row and j^th^ column.

UniFrac methods, following convention, were normalized via rarefaction (depth details by study found in Table 1). No additional normalization scheme was applied to novel methods in the gemelli Qiime2 package, in line with the 2022 publication and Qiime2 procedures.

Qiime2’s diversity modules were used to execute Jaccard, Bray-Curtis, weighted UniFrac, and unweighted UniFrac transformations and subsequent Principal Coordinate Analysis (PCoA). Qiime2’s gemelli plugin was used to perform RPCA and phylo-RPCA [17]. The gemelli plugin automatically performs an RPCA dimension reduction down to three principle components, so machine learning (ML) was initially conducted with three equivalent components for all beta diversity methods.

Following this, additional tables were created saving the top 3, 5, 10, 20, 50, and all MDS axes for Jaccard, Bray-Curtis, and UniFrac methods. These sets were used downstream to compare performance of different beta transformations with varying degrees of information. Due to the nature of gemelli plugin procedures, phylo-RPCA and RPCA remained static at 3 axes. Whole data and log-normalized data without beta transformation were also retained for ML.

### Statistics and Machine Learning

Univariate PERMANOVA was conducted using the R vegan package [18] with 999 iterations for each beta diversity distance matrix, once for each metadata column, following the general formula:

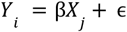

Such that *Y*_*i*_ is a distance matrix of the *i* ^*th*^ beta diversity approach and *X*_*j*_ is the *j*^*th*^ metadata feature from the provided studies.

Random forest and KNN were each used to relate beta diversity-associated distance matrices back to phenotype using the Python ML module, Scikit-learn[19]. Because RPCA-based methods only output 3 principle components, KNN and random forest were only performed on the first 3 axes for these methods. For all other beta diversity methods, ML models were run with the top 3, 5, 10, 20, and 50 MDS axes as input. ML models were also run once for each of the non-RPCA approaches with their full raw and log-normalized distance matrices.

Random forest was conducted 100 times with a stratified 0.4/0.6 test/train split for each qualifying metadata feature. Metadata features were filtered out if the test group size was smaller than the number of categorical classes available. Per categorical metadata feature, samples with unique classes that could not be trained or tested were also removed. For categorical features a classifier was used, while a regressor was used for continuous features.

The same quantitative filters were used in the KNN ML method. For categorical features, ROC-AUC values for each ML run were stored for comparison across metrics. For continuous features, *R* ^2^ was stored for each ML run.

Kruskal-Wallis testing was performed on *R*^2^ and ROC scores, with repeat values per ML run per feature across all datasets, grouped by beta diversity transformation, and subsequent Dunn tests were conducted with FDR correction for significant *R*^2^ and ROC scores.

### Script Validation and Black-Box Testing

To ensure reproducibility and reliability, black-box testing was conducted on core computational scripts used in the analysis. Testing scripts were designed to validate the original analyses by independently executing each script from start to finish using representative input data and producing comparable outputs for downstream use without altering the underlying functions of the original analysis. This black-box testing strategy emphasizes reproducibility by focusing on the validity of inputs and outputs rather than internal implementation details.

## Results

### Different Beta Diversity Transformations Appear to Resolve Phenotype with Similar ROC and R^2^ Values

In this study, we evaluated the impact of different approaches to data transformation and calculations of beta diversity on the ability of the random forest and KNN (Figure S1) algorithms to discriminate metadata features based on microbial relative abundance within five datasets. We used six beta diversity transformations, as well as the raw un-normalized and log-normalized counts. Following the procedure in a previous study [11], we assessed the first 3 EVD axes for ML for the six beta diversity metrics. Fig 1. summarizes the R^2^ values (for 36 quantitative variables) and ROC values (for 71 qualitative variables) across metadata features from all five datasets. Under this analysis scheme, the six beta diversity metrics show broadly similar performance across this wide analysis space (Figure 1). By comparison, inclusion of the raw and log-normalized count tables showed a clear increase in performance in comparison to all the beta diversity metrics for the quantitative variables (Fig. 1A); although this improvement in performance was not observed for the qualitative variables (Fig. 1B). We saw nearly identical results when utilizing KNN instead of random forest (sup. Fig 1) for ML. In general, there were no significant differences between the performance of the six ML algorithms, with the exception of weighted UniFrac which was slightly, but significantly, worse than the other beta diversity algorithms. Utilizing log-normalized and raw counts for ML yielded a significant improvement for the quantitative variables in comparison to the first 3 EVD axes but no significant difference for the qualitative variables.

**Figure 1:**
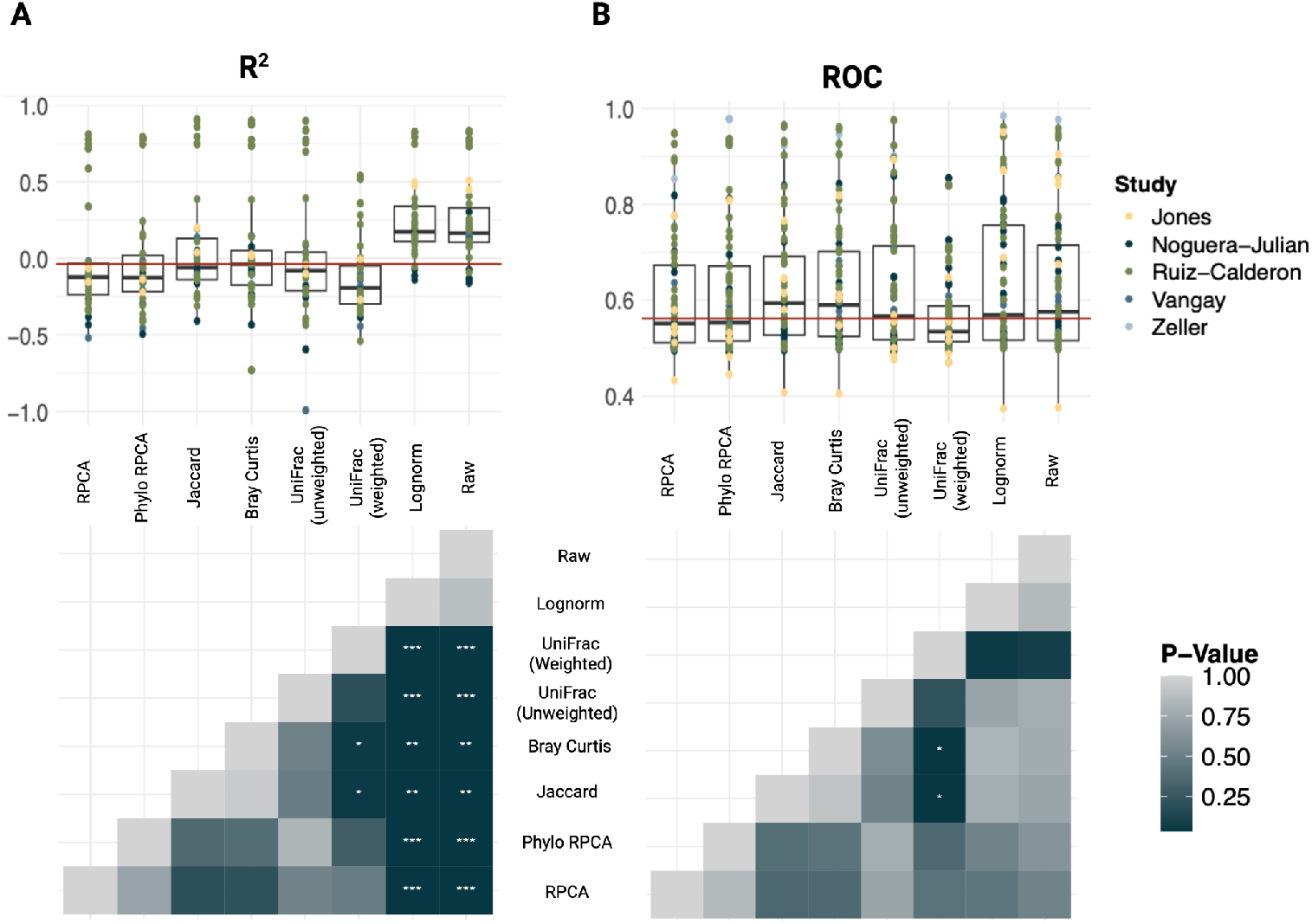
**(A)** Average random forest *R*^2^ values for each quantitative feature colored by study are shown across the six beta diversity metrics (top), where raw and log-transformed whole tables outperformed methods of beta transformation when resolving phenotype. This performance difference is confirmed by FDR corrected Dunn tests (bottom). **(B)** Average random forest for qualitative ROC values. There were no significant differences in phenotypic resolution, except an underperformance of weighted UniFrac compared to naïve Jaccard and Bray-Curtis methods, also seen in **(A)**.

### Inclusion of More EVD Axes in Random Forest Models Boost Naïve Performance Advantages, but Only Up to a Certain Point

The differences we observed between the beta diversity algorithms and the log-normalized and raw counts for quantitative variables could potentially be explained by two non-mutually exclusive hypotheses. These differences might have arisen from the properties of the beta diversity transformations. Or the ML algorithms might make use of all of the data represented in the log-normalized and raw count tables that are not present in the compressed representation of the data captured in only the first 3 EVD axes. To begin to discriminate between these two hypotheses, we ran the same random forest models, but incrementally increased the number of EVD axes we included for all applicable methods. We ran the beta diversity methods with three, five, ten, twenty, and fifty PCoA axes, as well as with the full EVD/PCoA matrix. RPCA and phylo-RPCA are only output by Qiime2 with 3 EVD axes, so they were not run with differing axis counts.

We found that the inclusion of more data in the form of increased number of principal components improved beta diversity performance as a feature predictor up through about the first 10 PCA axes (Fig. 2). Interestingly, including more axes, or all of the data, generated slightly decreased performance for all beta diversity metrics where these axes were applied. We conclude that EVD can be useful as a pre-processing filter, but the inclusion of only 3 EVD axes, as proposed by Martino et al. [11], may be too restrictive.

**Figure 2.**
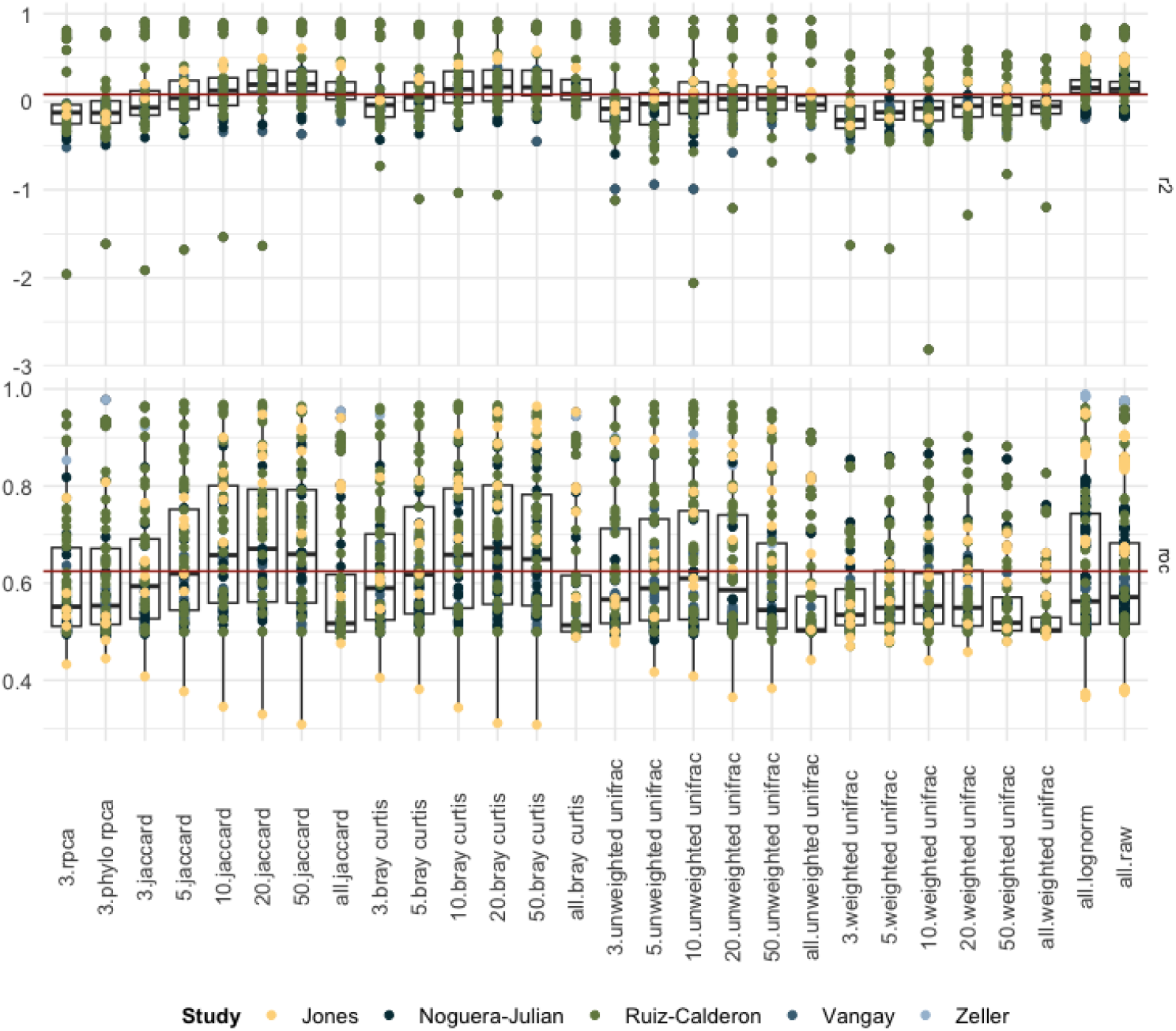
Average ROC and R^2^ values (Y-axes) for each metadata feature were plotted across beta diversity metrics where 3, 5, 10, 20, 50, and all MDS axes of the PCoA transformed tables were sent though ML. Increasing the amount of information provided to a random forest analysis shows an increase in all performance metrics, approaching the whole data (lognorm and raw) approaches up until about 20 axes. RPCA based methods are capped at 3 axes due to their standard output in Qiime2.

### Discrepancies Between Pseudo-F Values and P-Values in PERMANOVA May Cause Skewed Interpretations of RPCA-Based Transformations

In their paper, Martino et al. noted that utilization of the RPCA-based methods results in a large increase in pseudo-F values. Because we did not observe a corresponding increase in ML performance when using these algorithms in our analysis, we next examined the pseudo-F values they produced. Fig 3A shows the pseudo-F statistics from all six beta diversity metrics across the five datasets. We were able to successfully reproduce the observation from Martino et al. [11] that phylo-RPCA and RPCA have a much higher F-statistic for many of the metadata features. However, this increase was not associated with an increase in power for ML algorithms (Fig. 1). To explore this discrepancy, we ran 15 permutations on all the datasets, in which we shuffled the labels of all the data. Overall, there was little difference in the mean pseudo-F value across all permutations, but we noted an increase in variance in the pseudo-F statistic for phylo-RPCA and RPCA (Fig. 3B). This increase in variance, however, is not reflected in the p-values, which remain strictly uniform, as expected for our null shuffles across all six algorithms (Fig 3C). We conclude that an increase in variance of the pseudo-F statistic that are inherently associated with the RPCA based methods can at times lead to observations of larger pseudo-F statistics, but this more variable pseudo-F statistic does not seem to provide increased resolution for ML algorithms. Note that Figs 3A and 3B have extreme upper-end outliers removed from the phylo-RPCA methods for visualization purposes; this does not change the interpretation of results. See supplementary figure S2 for un-altered results.

**Figure 3.**
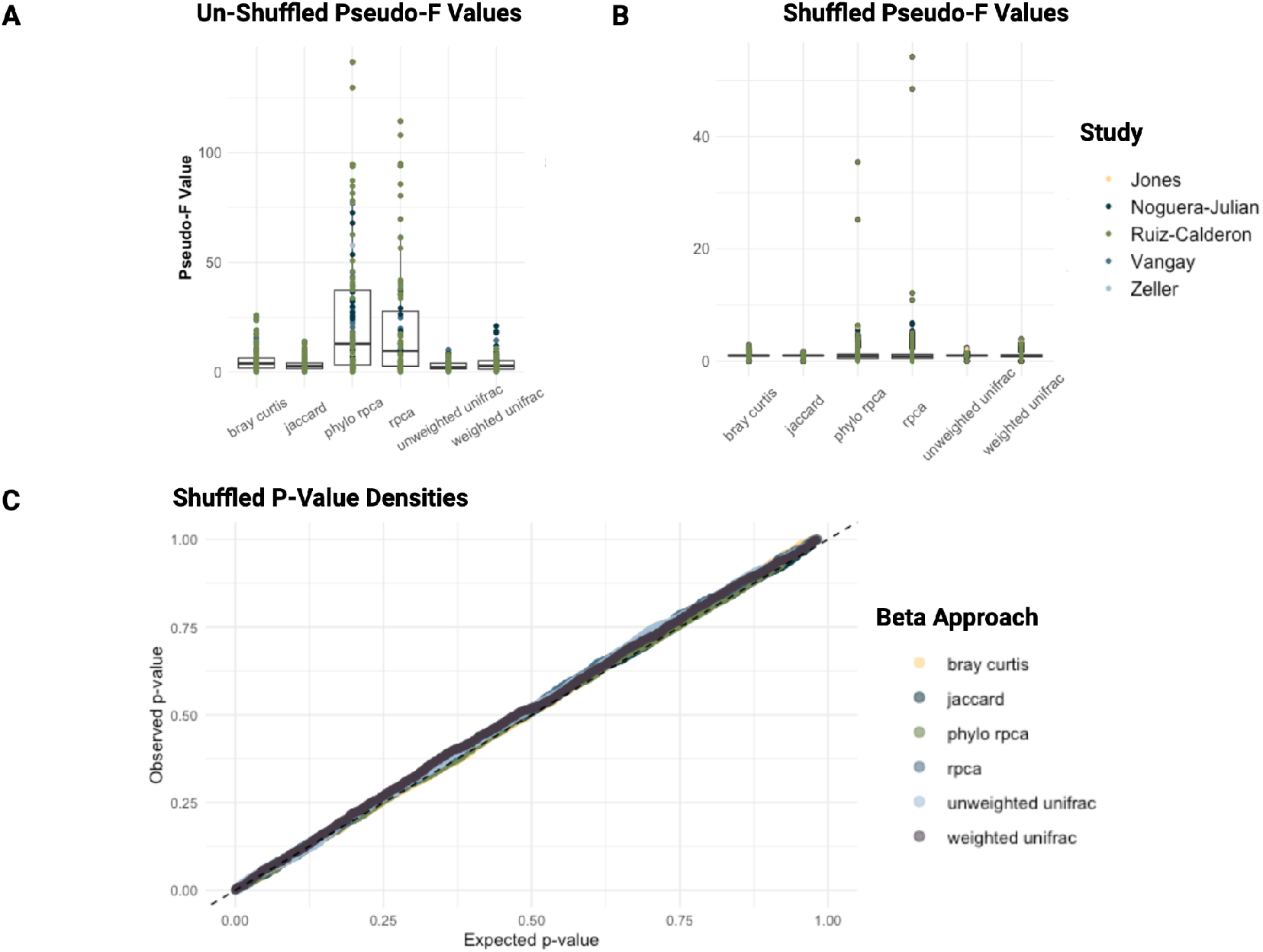
Comparison of PERMANOVA results under unshuffled and shuffled metadata conditions. **(A)** Unshuffled pseudo-F values for each study. **(B)** Shuffled pseudo-F values for each study, corresponding to the uniform distribution of shuffled p-values shown in panel C. **(C)** QQ-style plot showing expected p-values under a uniform distribution versus observed p-values from PERMANOVA runs with shuffled metadata labels. While shuffled p-values display the expected uniform distribution, several beta diversity approaches exhibit inflated pseudo-F values even after label shuffling, indicating unexplained deviations from randomized behavior.

## Conclusion

In their intriguing paper, Martino et al. promoted a novel pipeline with correction for sparsity and compositionality and evaluated this pipeline with K-nearest neighbor ML on the first 3 EVD axes. They reported this pipeline to be highly effective. Their evaluation was executed on two features within real datasets and a simulated dataset[11,17]. Our study expands on their observations in a number of important ways: (1) We evaluated five datasets (Table 1), including one of the datasets examined by Martino et al.; our analysis included 36 quantitative and 71 categorical variables, (2) We examined the use of random forest ML algorithms which represent algorithms that are much more widely used in the metagenomics literature than KNN, (3) We expanded our analysis beyond the 3 EVD axes, (4) We included pipelines that did not perform EVD prior to ML but rather used log-normalized or raw counts. (5) We examined the distribution of F-statistics generated by PERMANOVA in both the real data and shuffled datasets, in which we permuted the associated metadata. This approach follows the trends of recent papers suggesting that often simpler algorithms have equivalent or slightly improved performance over more sophisticated but also more complex approaches [10,20,21].

Upon completion of our own pipeline, we arrived at several key conclusions. Across datasets, there is no substantial difference between different beta diversity transformations when utilizing the first 3 PCoA axes for ML, with the exception of a slight decrease in performance for weighted UniFrac (Fig. 1). We obtained similar results from random forest and KNN models, indicating that choice of ML algorithm does not substantially impact the outcomes. Especially for quantitative variables, feeding raw or log-normalized counts to ML algorithms yields a notable increase in performance when compared to the beta diversity transformed data (Fig. 1A and 1B). When expanding beyond the first 3 EVD axes, ML algorithms show increased performance up through the first 10-20 axes but then show slightly decreased performance when adding additional EVD axes (Fig. 2). Feeding the first 10-20 axes into ML algorithms also yields slightly better performance than using raw or log-normalized data. It will be of interest to see if ML models based on artificial neural networks, which have an embedding layer that can also reduce the dimensionality of complex datasets, will be sensitive to the number of input axes or can perform equally well when presented with full count tables of log-normalized or raw data.

In reaching the above conclusions, we evaluated six beta diversity transformations for their ability to provide EVD axes that can be utilized by ML models to predict phenotype from microbial composition across five 16S rRNA datasets. When EVD was applied, all six transformations performed comparably with no significant difference in phenotype prediction, with the exception of weighted UniFrac which showed a modest but statistically significant decrease in performance relative to the naïve methods. This reduction may stem from its uniform application across datasets and features without parameter tuning in our approach, underscoring that some transformations may not generalize well across diverse ecological contexts. We posit that the average user of these methods may not be inclined to fine-tune parameters, opting instead for out-of-the-box defaults, making this approach worth studying.

One of the observations made by Martino et al. was that phylogenetically and compositionally aware metrics produced F-statistics in PERMANOVA that were larger in magnitude than traditional beta diversity approaches. We observed that pseudo-F statistics exhibited a broader range of values under RPCA-based approaches than under other transformations, with an overall higher mean pseudo-F statistic. Strikingly, though, this pattern persisted even after metadata labels were shuffled (Fig. 3). In these shuffled datasets, the PERMANVOA p-values followed the expected uniform distribution for null datasets despite the broader range of F-statistics. We conclude that elevated pseudo-F statistics do not reliably correspond to genuine increases in statistical significance and power. Pseudo-F statistics should therefore not be generally taken to quantify effect size and should be interpreted with caution when used as an evaluation of relative performance of beta diversity algorithms.

Taken together, these findings indicate that for phenotype resolution tasks, complex corrections for compositionality or phylogeny do not consistently outperform simpler, more direct approaches. Our analysis reflects a broad and minimally filtered set of metadata variables with varying degrees of biological meaning to approximate real-world analytical conditions. While including variables of variable relevance might reduce overall predictive accuracy, this approach captures how ML is often applied in exploratory microbiome analyses and provides a fair assessment of method robustness.

Despite the importance of these findings, this meta-analysis has several limitations. Although we expanded our evaluation beyond three eigenvalue decomposition (EVD) axes in some analyses, RPCA-based transformations were ultimately compared using only the first three axes based on Qiime2 output. This constraint may underrepresent the potential information retained in higher-dimensional spaces. Our decision to include a broad range of metadata variables also likely introduced additional variability into model performance. While this reflects realistic analytical conditions, it could potentially obscure differences among methods that may exist with more carefully selected features.

Our study also offers several strengths that enhance the robustness and interpretability of its findings. It leverages a comparatively large and diverse analytical framework including five independent 16S rRNA datasets and 107 metadata features. This allows for a more general assessment of method performance than prior studies limited to smaller or simulated datasets. We also implemented a consistent and uniform analytical pipeline across all datasets, minimizing confounding effects introduced by inconsistent preprocessing or tuning. This approach provides a realistic reflection of how these tools are typically applied in practice, especially by users employing default parameters rather than extensive optimization.

Ultimately, our results suggest that continued improvements in linking microbiome composition to phenotype might depend less on increasingly complex diversity transformations and more on complementary strategies: biologically informed feature selection, rigorous statistical validation, and integration with modern ML frameworks. These insights build on and expand the observations of Martino et al., demonstrating that while transformations that account for sparsity and compositionality offer theoretical appeal, their practical benefits in phenotype prediction remain limited under typical analytical conditions.

## Acknowledgements

We thank Megan Hill for helpful comments on a previous draft of this paper. This work was supported primarily by the Engineering Research Centers Program of the National Science Foundation under NSF Cooperative Agreement No. EEC-2133504. This work was supported by the University of North Carolina at Charlotte’s Department of Bioinformatics and Genomics.

## Supplemental Figures

**Figure S1:**
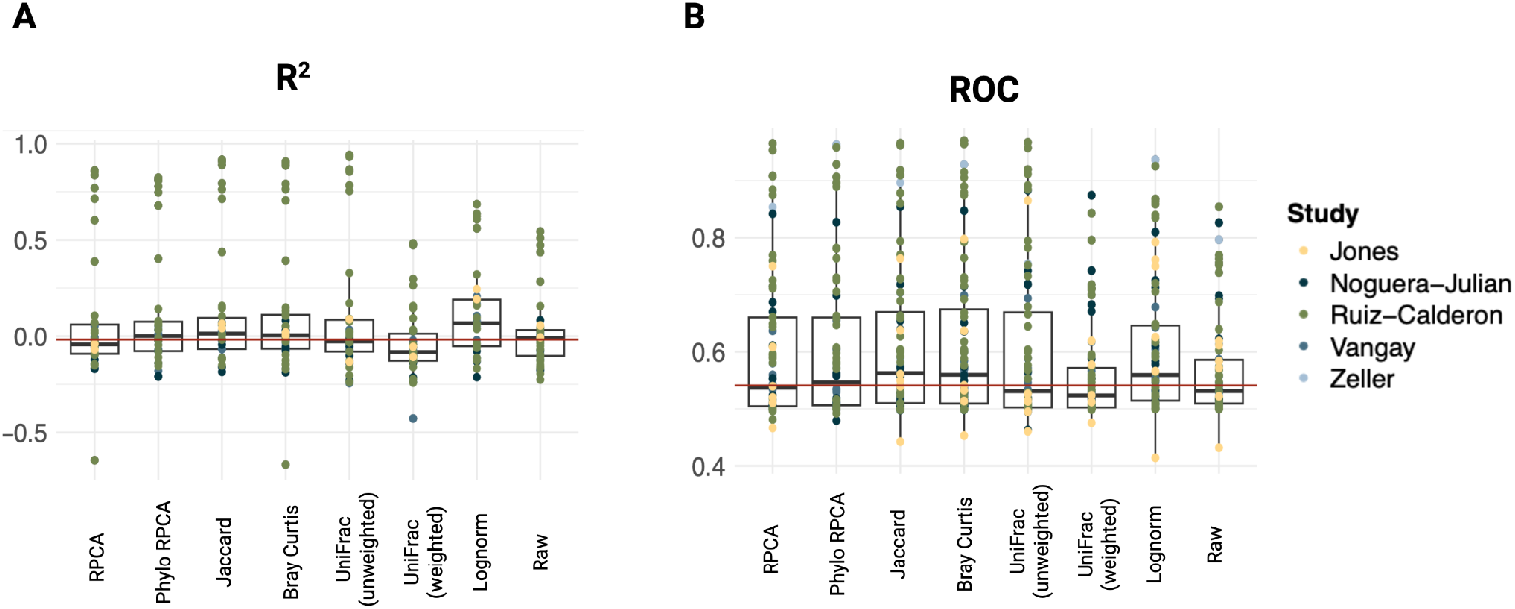
**(A)** Average KNN *R*^2^ values for each feature colored by study are shown across the 6 beta diversity metrics, where log transformed whole tables outperform methods of beta transformation when resolving phenotype. **(B)** Average KNN roc values, as with random forest, yield overall more subtle results.

**Figure S2.**
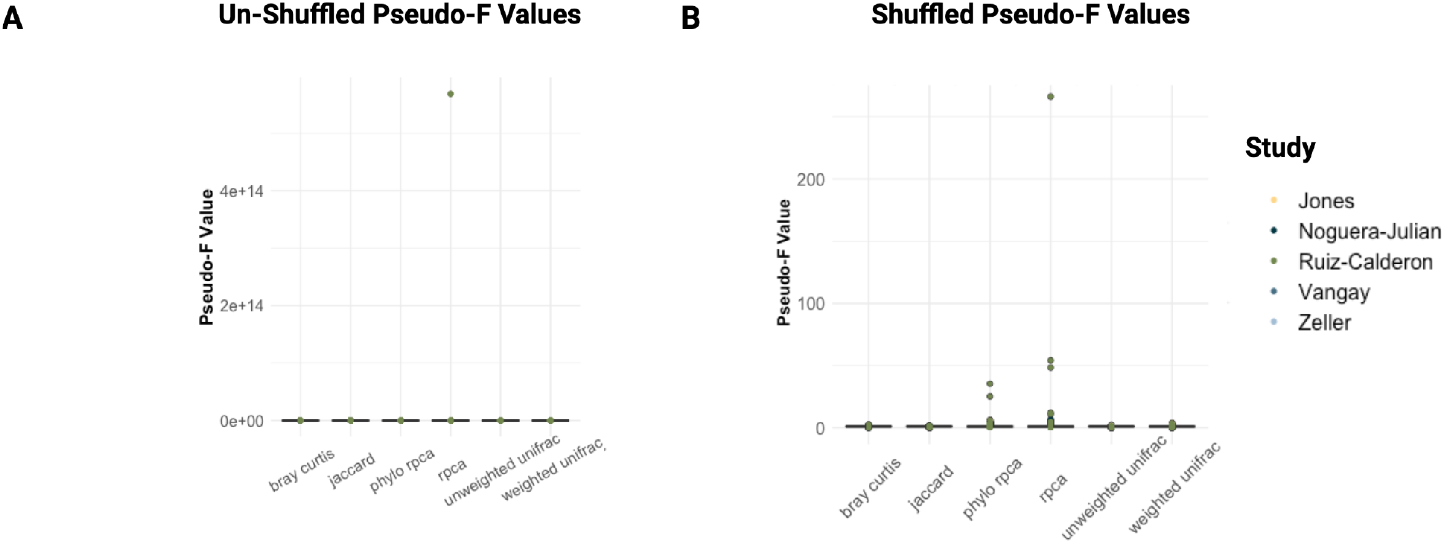
Comparison of PERMANOVA results under unshuffled and shuffled metadata conditions. **(A)** Unshuffled pseudo-F values for each study. **(B)** Shuffled pseudo-F values for each study.

## Notes

https://github.com/DaisyBrumit/beta_diversity_testing

### Competing Interest Statement

The authors have declared no competing interest.

